# Minimizing detection bias of somatic mutations in a highly heterozygous oak genome

**DOI:** 10.1101/2025.02.13.638107

**Authors:** Wenfei Xian, Pablo Carbonell-Bejerano, Fernando A. Rabanal, Ilja Bezrukov, Philippe Reymond, Detlef Weigel

## Abstract

Somatic mutations are particularly relevant for long-lived organisms. Sources of somatic mutations include imperfect DNA repair, replication errors, and exogenous damage such as ultraviolet radiation. A previous study estimated a surprisingly low number of somatic mutations in a 234-year-old individual of the pedunculate oak (*Quercus robur*), known as the Napoleon Oak. It has been suggested that the true number of somatic mutations was underestimated due to gaps in the reference genome and too conservative filtering of potential mutations. We therefore generated new high-fidelity long-read data for the Napoleon Oak (*n* = 12) to produce both a pseudo-haploid genome assembly and a partially phased diploid assembly. The high heterozygosity allowed for complete reconstruction of phased and gapless centromeres for 22 of the 24 chromosomes. On the other hand, the high heterozygosity posed challenges for short-read alignments. Use of only the pseudo-haploid assembly as a reference led to potential misalignments, while use of only the diploid assembly reduced variant detection sensitivity. Since most somatic mutations are layer-specific, the observed frequency is expected relatively low, even where all cells in a single layer contain a specific mutation. To address this challenge, we employed a read assignment strategy, selecting the appropriate reference sequence (pseudo-haploid or diploid) based on alignment score and mapping quality. Ultimately, we identified 198 high-confidence somatic mutations, compared with 17 somatic mutations identified before with the same set of short reads. Our approach thus increased the total estimated annual mutation rate by a factor of five.

## Introduction

Fifteen years ago, a seminal study by Sally Otto and colleagues showed that mutations reducing male fertility accumulate in a long-lived deciduous tree in a clock-like manner, suggesting that there are natural limits to how long trees could produce viable offspring (Ally, Ritland and Otto, 2010). Since that early paper, a series of studies has attempted to directly detect the accumulation of mutations in trees using a range of sequencing strategies, resulting in a remarkably wide range of mutation rate estimates (summarized in (Johannes, 2024)).

Oaks, widely distributed in Northern Hemisphere forests, are renowned for their longevity, with many individuals living for centuries (Leroy, Plomion and Kremer, 2020). One noteworthy specimen, known as the Napoleon Oak, which grows on the campus of the University of Lausanne, is over 200 years old and was the subject of a recent somatic mutation study (Schmid-Siegert *et al*., 2017), which used a reference genome assembled from long reads of this individual in combination with short reads of material from a lower and an upper branch. Ten single nucleotide variants (SNVs) were identified in the upper branch - sample 66 and seven SNVs in the lower branch - sample 0 (Schmid-Siegert *et al*., 2017). Subsequent work with another oak tree suggested that the number of somatic mutations identified in the Napoleon Oak was a likely underestimate because the minimization of false positives (specificity) had been prioritized over the identification of all true positives (sensitivity). Apart from the reference genome having been generated with an early long-read technology, the threshold for variant calling might have been overly stringent (Plomion *et al*., 2018). In addition, the high heterozygosity of oak could have impacted variant calling, as only a single phase of the genome had been assembled.

When reads are aligned to a haploid assembly, regions with high sequence diversity between the two haplotypes might cause misalignments, as a haploid reference cannot fully represent both haplotypes. This has been quantitatively confirmed in highly heterozygous sweet oranges, where a phased assembly as reference yielded more than twice the number of somatic mutations compared to a haploid assembly (Wang *et al*., 2024). Using a diploid reference is, however, not without its own challenges. In regions that are identical in the two haplotypes, aligners will randomly assign reads to one of the two phases, reducing the power of variant identification, particularly when there are only very few variant reads covering a specific position. Conversely, high-frequency variants may appear on both haplotypes, leading to an overestimate of true variants.

A recent study of two tropical trees found that almost all somatic mutations are present at low frequency in the sampled tissue (Schmitt *et al*., 2024). Other studies have confirmed that most somatic mutations are layer-specific, as has long been known from the study of mutations with phenotypic impact, such as grape color (Dermen, 1960; Amundson *et al*., 2023; Goel *et al*., 2024). Since the samples sequenced in the Napoleon Oak study were entire leaves (Schmid-Siegert *et al*., 2017), this might have further reduced the observed frequency of detected somatic mutations.

Here, we produced a new genome assembly of the Napoleon Oak and optimized a somatic mutation calling pipeline for highly heterozygous genomes, using both haploid and diploid reference genomes. Utilizing this improved approach, we identify a larger set of high-confidence somatic mutations in the Napoleon Oak, which lead to a substantial upward revision of the annual somatic mutation rate.

## Results

### Genome Architecture of *Quercus robur*

To identify high-quality somatic mutations in the diploid pedunculate oak (*Quercus robur, n* = 12) individual known as the Napoleon Oak, we first assembled a high-quality genome of this individual. In August 2021, fresh leaves were sampled from the bottom branch, at the same location where material used for genome assembly and mutation calling (sample 0) had been sampled before (Schmid-Siegert *et al*., 2017). Two PacBio SMRTcells were used to generate a total of 63.7 Gb of HiFi data (**Table S1**). Using Hifiasm (Cheng *et al*., 2021), we assembled a highly contiguous haploid genome comprising only 16 contigs (**Figure 1A**).

**Figure 1.**
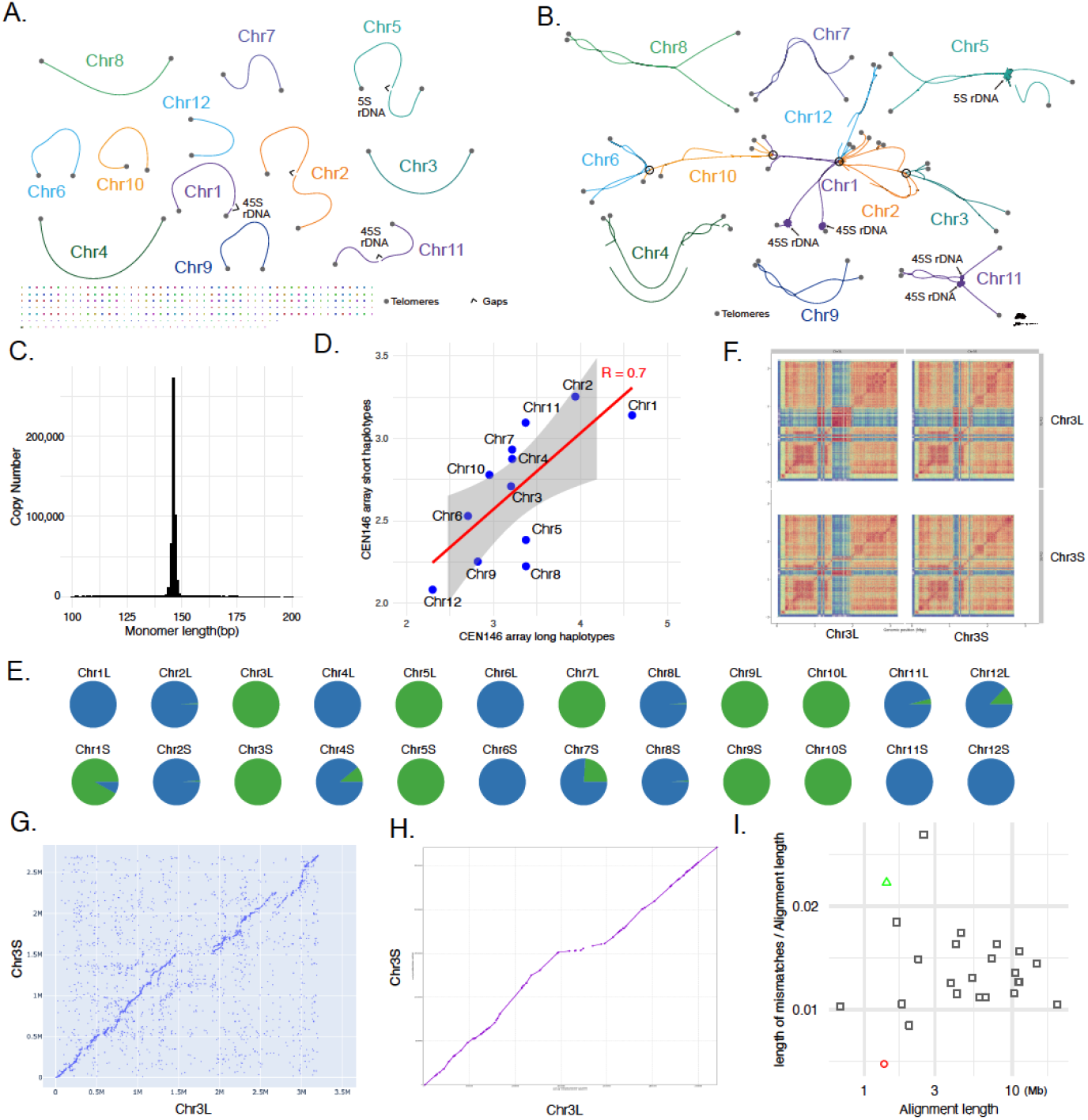
Genome Architecture of *Quercus robur*. A. Scaffolded haploid assembly graph. B. Contig-level diploid assembly graph. C. Copy number of satellite repeats in the diploid assembly. D. Length of the CEN146 array. E. Orientation distribution of CEN146: forward strand (green) and reverse strand (blue) proportions for each chromosome (columns) and haplotype (rows). F. Heat maps depicting sequence identity within and between the two haplotypes of the CEN146 array on Chr3. G. Dot plot of rare kmers showing that the two haplotypes of the CEN146 array on Chr3 can be aligned with Unialigner. H. Minimap2 alignment of the CEN146 array on Chr3. I. Substitution rates across various regions: green triangle represents Chr3 CEN146 array alignment using Minimap2, red circle indicates alignments using Unialigner, and gray rectangles indicate unique non-CEN146 sequences aligned with Minimap2. The details of the unique regions were recorded in Table S7.

Eight of the contigs contained telomeric repeats at both ends, and thus corresponded to eight chromosomes. Due to the high heterozygosity of oak, the k-mer-based heterozygosity estimate is about 1.6% (**Figure S1**). As a result, the haploid assembly cannot represent the entire genome. Therefore, we assembled a diploid genome from the same HiFi data using Verkko (Rautiainen *et al*., 2023) (**Figure 1B**).

The assembly of plant rDNA clusters remains a challenge (Rabanal *et al*., 2022; Fultz *et al*., 2023). The genome of the pedunculate oak contains two 45S rDNA clusters and one 5S rDNA cluster (Bočkor *et al*., 2014). We used information from the Verkko assembly graph to scaffold the remaining eight contigs of the Hifiasm assembly into four chromosomes (**Table S2**). In the Verkko assembly graph, rDNA clusters formed tangles and connected sequences at both ends (Rautiainen *et al*., 2023). We aligned the eight contigs of the Hifiasm assembly without telomeres on both ends to the diploid assembly and projected them on the diploid assembly graph (**Figure S2**). Based on the connections in the graph, we manually scaffolded the eight contigs, resulting in four additional chromosome-level pseudomolecules. Each had one end capped by telomere repeats and large arrays of tandem repeats (either rDNA or another, unknown type) at the other end (**Table S2**). The final haploid genome had high accuracy (QV 53) and high completeness (BUSCO 99.8%), significantly surpassing the quality of the previous version generated from an earlier generation of long reads (**Figure S3** and **Table S3**) (Schmid-Siegert *et al*., 2017). The assembled haploid genome spans 810 Mb, in agreement with estimates for the 1C genome size for this species, which range from 759 Mb to 1,068 Mb (*Plant DNA C-values Database*, no date).

The long and highly accurate PacBio HiFi reads enabled the assembly of repetitive regions, such as centromeres (Naish *et al*., 2021). In each of the 12 chromosomes of the haploid assembly, we identified large arrays of a shared 146 bp satellite repeat unit, with a genome-wide total of 486,051 copies in the diploid assembly (**Figure 1C, Figure S4** and **Table S4**), with high levels of CpG methylation (**Figure S5**). These arrays, on average about 3 Mb long, are likely centromeres, of which 22 of the 24 were gapless and phased. The CEN146 arrays on chromosome two were manually scaffolded (**Table S5**). Uniform coverage of HiFi reads alignment in these regions indicates no significant assembly error (**Figure S6**). Within the CEN146 arrays, transposons were rare but transposons can be observed in the flanking regions (**Figure S6**). We categorized the CEN146 arrays of homologous chromosomes into long haplotype (L) and short haplotype (S) based on their lengths. The size of CEN146 arrays differed more between chromosomes than between haplotypes of each chromosome (**Figure 1D**). The relative orientation of CEN146 repeats within the arrays were haplotype- rather than chromosome-specific (**Figure 1E**). For instance, on Chr1S, Chr4S, Chr7S and Chr12L, more than 8% of repeat units were in opposite orientation, whereas their counterparts (Chr1L, Chr4L, Chr7L and Chr12S) had less than 1% of arrays in the opposite orientation.

Due to the challenges of aligning long highly repeated sequences, previous studies have often greatly differed in their estimates for evolutionary rates of centromeres (Logsdon *et al*., 2021; Bzikadze and Pevzner, 2023). Here, we attempted to use our phased CEN146 arrays to examine the two alignment approaches: Minimap2 (Li, 2018), based on standard concepts of molecular evolution, and Unialigner (Bzikadze and Pevzner, 2023), which is designed for aligning long tandem repeats. First, we used Unialigner to align the two haplotypes of the CEN146 arrays of each chromosome. Dot plots of rare kmers revealed that only chromosome 3 contained a substantial number of rare kmers, indicating that alignment was feasible only for chromosome 3 (**Figure 1G** and **Figure S7**). Because only chromosome 3 produced a Minimap2 alignment consistent with the Unialigner dot plot (**Figure 1H** and **Figure S8**), we focused on comparing the CEN146 arrays of chromosome 3 (**Figure 1F**).

Since Unialigner uses rare kmers as anchors to extend alignments, we used the alignment blocks of Minimap2 with the maximally possible mapping quality (MAPQ) of 60. The total alignment lengths were similar for the two tools, with Unialigner aligning 1.36 Mb and Minimap2 aligning 1.41 Mb. However, the substitution rate inferred by Unialigner is 0.004, which is five times lower than that of Minimap2, 0.02. Notably, the substitution rate from Minimap2 was even higher than that of most non-CEN146 uniquely aligned regions using Minimap2 (**Figure 1I** and **Table S6 & 7**). Because the two tools yield such diverging results, we did not consider the centromeres for our estimates of somatic mutation rates either.

### Variant Allele Frequency Estimates of Layer-Specific Mutations

Our goal is to identify somatic mutations that had become fixed in stem cells that gave rise to the sampled tissue (Schmid-Siegert *et al*., 2017). We take the cell layer where the mutations occur into account, since the layer specificity of somatic mutations has not only long been known from phenotypic examination (Dermen, 1960), subsequently confirmed by molecular analysis of specific mutations (Kobayashi, Goto-Yamamoto and Hirochika, 2004), but has also recently shown to be the prevailing type by high-throughput sequencing in potato and apricot tree (Amundson *et al*., 2023; Goel *et al*., 2024).

The plant body comprises multiple, segregated layers of cells that arise from multiple layers of stem cells in the meristems (Steeves and Sussex, 1989), where the outermost layer, L1, makes up the epidermis, the L2 the photosynthetic tissue, and the L3 the ground tissue. The frequency with which fixed somatic mutations are observed in a sample that includes all layers will depend on the relative contribution of each layer to the total genomic DNA in that sample. Although we lack layer-specific data to evaluate the DNA content of each layer in oak leaves, one can roughly approximate the DNA content of the L2 by analyzing shared mutations detected in leaves and acorns, since embryonic tissues originate from the L2 (Amundson *et al*., 2023).

In the published oak accession 3P, nine somatic SNVs were shared between leaves and the embryonic tissue in acorns (Plomion *et al*., 2018). Thus, these nine SNVs are most likely derived from the L2 of the meristem and can be considered as L2 markers. The median frequency of reads covering these nine somatic mutations in the leaves was approximately 0.3 (**Table S8**). Since a fixed mutation will appear in only one haplotype, the DNA of the mutant cells would have constituted 60% of the total DNA of the sample (**Figure 2A**). This provides an upper bound for the contribution of the other two layers. Since the L1 contributes the least amount of DNA, we infer that L1 < L3. Given that L1 and L3 together make up about 40% of all reads, this would suggest that the L1 contributes <20% and L3 >20%. The expected observed frequency of L1-fixed SNVs in the haploid assembly, which serves as the reference, would thus be below 0.1 (below half of 20%). Therefore, the use of a higher variant allele frequency (VAF) threshold may prevent the identification of fixed somatic mutations specific to certain layers.

**Figure 2.**
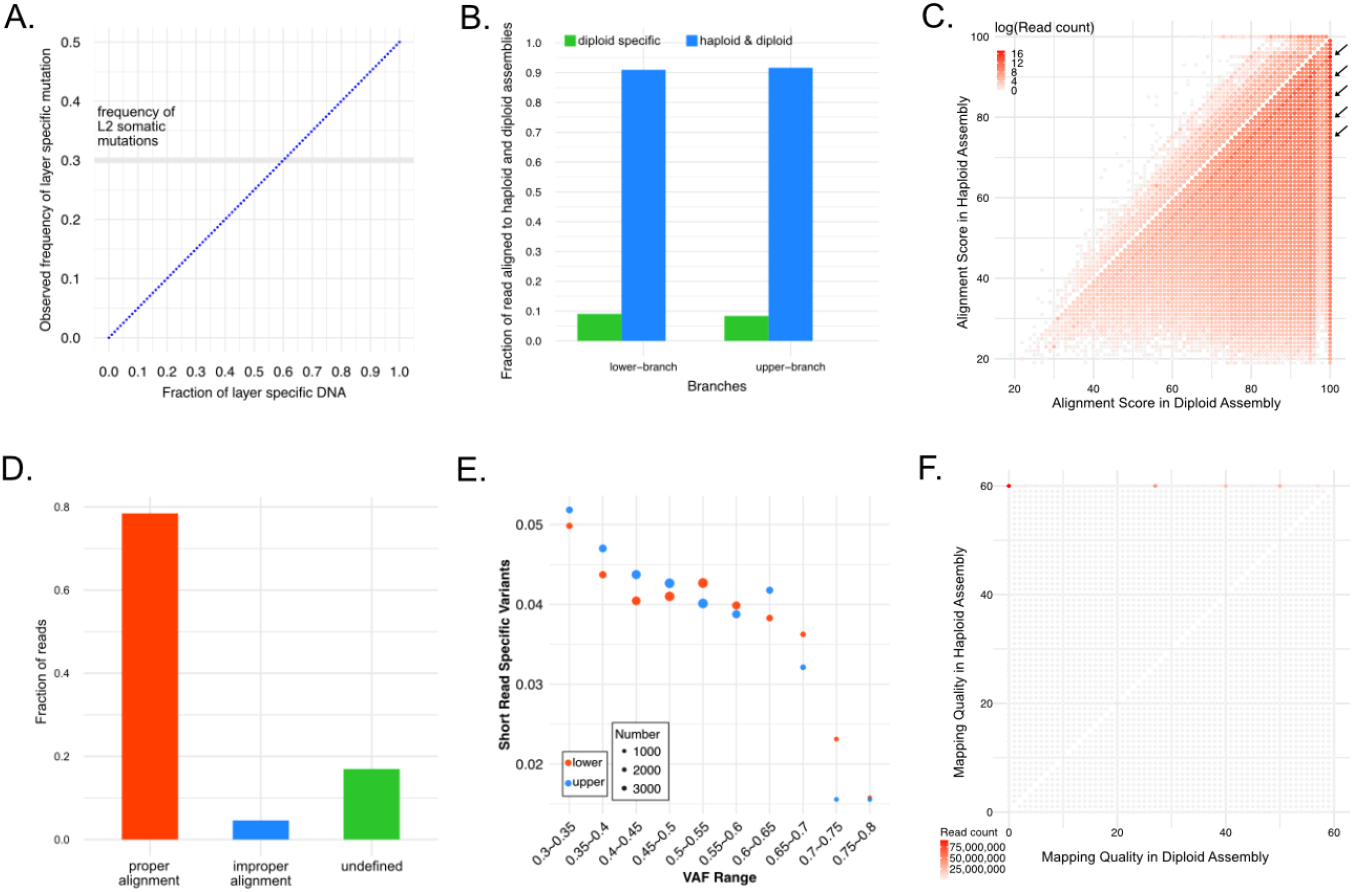
Using haploid or diploid assemblies for short-read alignment. A. Simulation of observed frequencies of somatic mutations relating to the contribution of a layer to the total DNA of the sample. B. Fractions of reads aligned to only the diploid assembly. C. Distribution of alignment scores in two assemblies as reference. The dots indicated by arrows represent the germline single nucleotide variants. D. Reads whose alignment position matched the expected position (as inferred from alignments of 4.1 kb diploid assembly fragments centered on each read to the haploid assembly) were classified as having a proper alignment. Reads that aligned outside the expected position were classified as having an improper alignment. Undefined are reads where the 4.1 kb diploid assembly region centered on the read aligned to a region in the haploid assembly that was smaller than 4 kb. E. Fraction of variants identified with short reads that were not identified from HiFi reads. The size of the dots indicates the number of short-read specific variants. F. Distribution of mapping qualities when using either the haploid or diploid assembly as reference.

### Limitations of Using Either a Single Haploid or Diploid Assembly as Reference

In highly heterozygous genomes, a single haploid assembly is an imperfect representation of the entire genome, and reads from regions that are highly diverged in the not represented haplophase cannot be aligned. We compared the short read IDs that mapped to the haploid assembly versus those mapped to the diploid assembly and found that approximately 9% of the reads could only be aligned to the diploid assembly (**Figure 2B, Table S11**). Therefore, using only the haploid assembly as the reference genome increases the false-negative rate in mutation identification.

Even though more reads can be mapped to the diploid assembly than to the haploid assembly, the number of paired reads with opposite orientation or mapping to different chromosomes is reduced (**Table S12**). Additionally, when reads are mapped to the diploid assembly, the average size of mapped fragments is smaller, with reduced standard deviation. This suggests that mapping to the diploid assembly can increase read mapping accuracy and reduce false positives.

We extracted reads that mapped to both assemblies and compared their alignment scores in the haploid and diploid assemblies (**Figure 2C, Figure S9**). Although a correct alignment generally results in a higher alignment score, when aligning short reads to a haploid genome, heterozygous sites appear as mismatches, with BWA assigning +1 for a match and -4 for a mismatch by default (Li and Durbin, 2010). For reads with an alignment score of 100 in the diploid assembly, alignment scores in haploid assembly values are predominantly 95, 90, 85, 80 or 75. Overall, aligning reads to the diploid genome increases the alignment score, indicating a reduction in alignment errors.

Because it is much easier to correctly align long sequences than it is to align short reads, we first aligned the short reads to the diploid assembly to serve as a reference framework and then extracted the flanking 2 kb sequences for every read and aligned them to the haploid assembly with minimap2. Using the flanking sequences from the diploid assembly as an intermediary, we inferred the expected positions of short reads in the haploid assembly and compared these to the actual alignment positions when short reads were mapped directly against the haploid genome. Reads that did not align within the range of expected positions were identified (**Figure S10**). Such improperly aligned reads accounted for 5% of the total reads mapped to the haploid assembly. (**Figure 2D, Table S14**).

Next, we investigated potential errors in the haploid assembly by identifying germline SNVs. We used DeepVariant (Poplin *et al*., 2018) to call germline SNVs on alignments of short reads from either the upper or lower branch to the haploid assembly. To ensure the reliability of variants identified in the short-read dataset, we retained only variants with a depth between 0.5x and 2x sample coverage, a minimum VAF of 0.3, and those marked as PASS. Because DeepVariant may produce false negatives, we extracted the variants identified in the short reads data and used Bam-readcount to verify whether these variants were present in the HiFi data. We applied a very lenient threshold: if a variant was supported by at least 5 HiFi reads (the genome wide coverage was 77x), we considered it reproduced in HiFi data. As expected, the fraction of short read-specific variants decreased as the VAF of short read variants increased. Notably, when the VAF of short read variants was between 0.3 and 0.35, the proportion of short read-specific variants remained around 5% (**Figure 2E, Table S13**).

While aligning reads to a diploid genome can mitigate alignment errors caused by differences between the two haplotypes, using only the diploid genome also has limitations. For instance, in homozygous regions, MAPQ is often 0 (due to multiple mapping), and coverage is reduced by half, making low-frequency mutations more difficult to detect. Among reads with different MAPQ values when mapped against either the haploid or diploid reference assembly, most had a MAPQ of 60 in the haploid assembly, but a MAPQ of 0 in the diploid assembly (**Figure 2F**).

### Optimizing Short-read Alignments in Highly Heterozygous Genomes via Dynamic Reference Genome Selection

To overcome these issues, we utilized both haploid and diploid assemblies as reference genomes. The haploid assembly was employed to identify variants in regions identical in both haplotypes, while the diploid assembly was leveraged to resolve heterozygous regions unique to each haplotype. We followed a hierarchical decision-making strategy, in which reads were first aligned to both haploid and diploid assemblies and each read was then assigned to one of the two reference assemblies based on a simple prioritization rule: If a read had a higher alignment score in one assembly, it was assigned to that assembly. If alignment scores were identical, the read was assigned to the assembly with the higher MAPQ value. Finally, if both alignment scores and MAPQ values were equal, the read was assigned to the diploid assembly (**Figure 3A**). The details of the variant calling pipeline and subsequent filtering criteria are described in the Methods.

**Figure 3.**
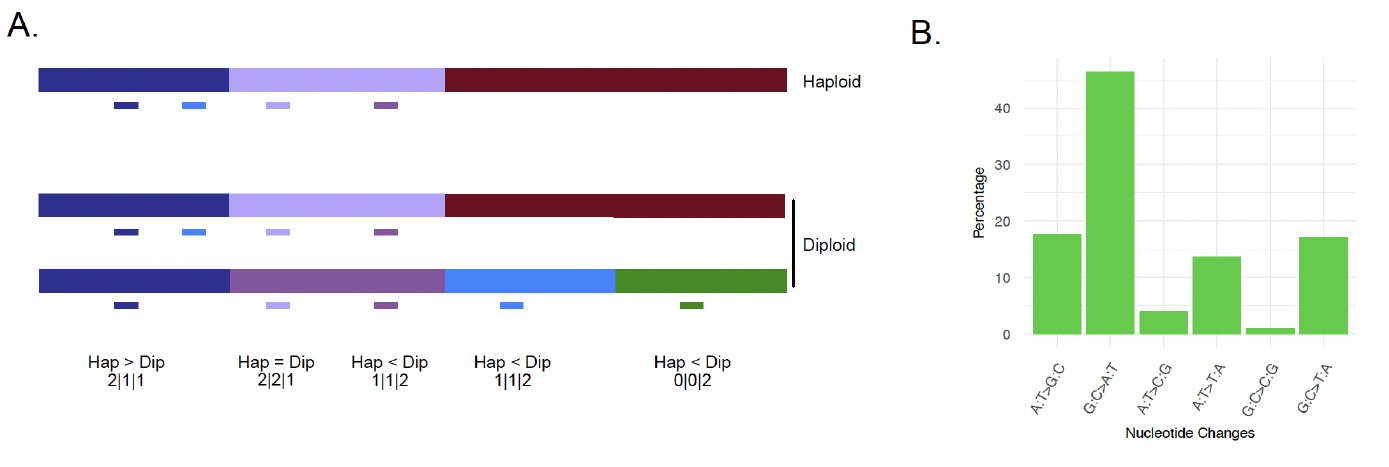
Somatic mutations in the Napoleon Oak. A. Diagram of how inappropriately aligned short reads in the haploid and diploid assembly can be identified. Scores indicate alignment in haploid and in the diploid genome composed of two haplophases. 2, high alignment and MAPQ scores. 1, high alignment but low MAPQ scores, or low alignment but high MAPQ scores. 0, no alignment. Dark blue read: homozygous region, reads in both assemblies have the same alignment scores, but MAPQ value is higher in haploid than in diploid assembly. Light purple read: heterozygous region, read has higher alignment score and MAPQ value in haploid versus diploid assembly and alignment score differs between haplophases. Dark purple read: heterozygous region, read has higher alignment score in diploid versus haploid assembly and alignment score differs between haplophases. Light blue read: heterozygous region, read has higher alignment score in diploid versus haploid assembly and alignment score differs between haplophases. Green read: highly divergent region, read only aligned to one haplophase of diploid assembly. B. Mutation spectrum of all 198 somatic SNVs.

### Somatic Mutations in the Napoleon Oak

Initially, with reads assigned to the diploid assembly as reference, 137 mutations were called in the upper branch and 44 mutations in the lower branch. From reads assigned to the haploid assembly as reference, 53 mutations were called in the upper branch and 21 mutations in the lower branch. We then visualized the results in the Integrative Genomics Viewer (IGV) for inspection (Robinson *et al*., 2017), including the separate BAM files for short reads from both branches, the original BAM file of short reads from both branches, and the BAM file of HiFi reads used to generate the assemblies. IGV screenshots of both retained and SNVs removed with our manual filters are shown in **Figure S11** (available on Figshare).

After IGV inspection, we retained 123 mutations in the upper branch and 14 in the lower branch with the diploid assembly as reference, and 51 mutations in the upper branch and 11 in the lower branch with the haploid assembly as reference. Combining the two datasets resulted in 173 mutations in the upper branch and 25 in the lower branch. Because we had called mutations in the upper branch using the lower branch as control and vice versa, there was by definition no overlap between the two branches, resulting in a total of 198 SNVs that were absent from the germline (**Table S9**).

All of the ten upper-branch variants that had been previously validated by amplicon sequencing (Schmid-Siegert *et al*., 2017) were identified with our approach: two were detected with the diploid assembly as reference, and eight were detected with the haploid assembly as reference (**Table S10**). As a control. We called SNVs using Strelka2 (Kim *et al*., 2018), a tool specifically developed for rare frequency somatic mutation. Of the ten validated upper-branch SNVs (Schmid-Siegert *et al*., 2017), Strelka2 identified nine SNVs with the haploid assembly as reference. One SNV could not be detected because its region of origin is absent from the haploid assembly. With the diploid assembly as reference, Strelka2 identified seven of the ten validated SNVs. Inspecting the alignments of the three missing SNVs in IGV indicates that failure to detect them with Streka2 is likely that the MAPQ values of the corresponding reads is 0 (**Figure S12**).

For the seven lower-branch variants confirmed by amplicon sequencing, we did find rare variant reads for these sites also in the upper branch, both in the original and filtered BAM files. Therefore, we do not consider these seven variants to be true lower-branch-specific mutations compared to the upper branch.

Consistent with mutation spectra observed in other systems (Hofmeister *et al*., 2017; Exposito-Alonso *et al*., 2018; Weng *et al*., 2019; Belfield *et al*., 2021; Satake *et al*., 2023), we found that G:C > A:T mutations were predominant (**Figure 3B**).

### Estimating Germline Mutation Rates

To approximate the per-site germline mutation rates in the Napoleon Oak, we considered only mutations that could be transmitted to the next generation, meaning they should have occurred in the L2. We reasoned above that the L2 accounts for around 60% of the total DNA in leaves. Each haploid set of chromosomes would thus contribute around 30% of the leaf DNA. To identify likely L2 mutations, we therefore used a VAF threshold of at least 0.2 for the haploid assembly as reference and 0.4 for the diploid assembly as reference. We found 77 such L2 mutations. For the Napoleon Oak, this would yield a raw mutation rate of approximately 6 × 10^−8^ bp^-1^. The annual mutation rate would be approximately 2.5 × 10^−10^ bp^-1^, within the range of rate estimates for other tree species (see the summary in (Johannes, 2024)). We do not address false negative rates, but even if we accepted all of our 198 originally called mutations to have been fixed in the L2 of the sampled leaves, the mutation rate estimate would still be within the range published for other trees.

For acorns produced by the Napoleon Oak today, which was 234 years old at the time of sampling (Schmid-Siegert *et al*., 2017), this would also be close to the generational mutation rate, since we are mostly ignoring mutations that occur since the entire branch was formed. Assuming that most oak trees have, however, only a generation time closer to 50 years, the generational mutation rate would have about 1.2 × 10^−8^ bp^-1^ as a lower bound. This is not very different from the per-generation mutation rate of *Arabidopsis thaliana* in nature, estimated to be about 3 × 10^−9^ bp^-1^ (Exposito-Alonso *et al*., 2018).

To assess the per-generation mutation rate, it is also useful to consider the number of cell divisions separating each generation. In the annual species *Arabidopsis thaliana*, it has been estimated that there are around 30 such cell divisions (Hoffman *et al*., 2004; Watson *et al*., 2016). In trees, there are perhaps twice to three times as many cell divisions separating each generation (Burian, Barbier de Reuille and Kuhlemeier, 2016). Thus, the per-site per-cell division mutation rate in oak and *A. thaliana* would be very similar. Note that we focus on mutations that are likely to be transmitted to the next generation, and that our study therefore does not speak to whether or not an allometric scaling law needs to be invoked for somatic mutation rates in plants (Johannes, 2024).

## Methods

### Biological Material and Pacbio HiFi Sequencing

Six grams of leaves were sampled on August 2021 from the bottom branch of the Napoleon Oak (*Q. robur*) on the campus of the University of Lausanne (Switzerland, 46° 31′ 18.9″ N, 6° 34′ 44.5″ E). Leaves were stored in liquid nitrogen. Prior to DNA extraction, to avoid contamination from pathogens, we removed any visibly spotted areas of the leaves. To improve DNA extraction efficiency, we also trimmed away the leaf veins. High molecular weight (HMW) DNA was obtained from frozen leaf powder as described by Calderon et al., 2024 (Calderón *et al*., 2024). In brief, the Nanobind Plant Nuclei Big DNA Kit (Circulomics) was used to extract HMW DNA from nuclei isolated according to Workman et al 2018 (*Website*, no date). We sheared the HMW DNA into appropriately sized fragments using a g-TUBE, then prepared a PacBio library using the SMRTbell Express Template Prep Kit 2.0 from 10 µg of sheared DNA. The library was selected for >10 kb fragments using a BluePippin instrument (Sage Science). Sequencing was performed using two 8M SMRTCells on a Sequel II.

### Genome Assembly and Annotation

PacBio CCS (*GitHub - PacificBiosciences/ccs: CCS: Generate Highly Accurate Single-Molecule Consensus Reads (HiFi Reads)*, no date) (v6.4.0) was used to generate CCS reads from the raw subreads with the parameter --min-rq=0.88. Subreads were aligned to CCS reads with ACTC (*GitHub - PacificBiosciences/actc: Align subreads to ccs reads*, no date) (v0.3.1). CCS reads were further polished by DeepConsensus (Baid *et al*., 2023) (v1.2.0) resulting in 63.65 Gb high-quality HiFi reads.

From cytology we know that pedunculate oak has two 45S rDNA clusters and one 5S rDNA cluster (Bočkor *et al*., 2014). Because current sequencing technologies still struggle to effectively resolve highly repetitive rDNA clusters (Rabanal *et al*., 2022). Thus, the ideal haploid telomere-to-telomere assembly would have only three gaps, resulting in 15 contigs. HiFi reads were assembled using Hifiasm v0.24.0 (Cheng *et al*., 2021), producing 16 long contigs. Based on length ranking, the 16^th^ contig was 3.5 Mb long, while the 17^th^ contig was much smaller, only 0.5 Mb. Therefore, we retained only the 16 longer contigs. Among these, eight contained telomeric repeats at both ends, and the remaining eight contigs had telomeric repeats at only one end.

Commonly used scaffolding methods rely on reference genomes (Alonge *et al*., 2022), but this approach may introduce reference bias. Moreover, only four gaps remained in our assembly, three of which likely contained rDNA clusters. Previous studies indicated that rDNA clusters form tangles in the Verkko assembly graph, and we visualized a Verkko v1.4.1 (Rautiainen *et al*., 2023) assembly with Bandage v0.9 (Wick *et al*., 2015) to identify tangles caused by rDNA clusters.

Using Minimap2 2.24-r1122 (Li, 2018), we aligned the eight contigs with telomeric repeats at only one end to the Verkko assembly. From the unitig-popped.layout.scfmap file, we identified the corresponding nodes in the assembly graph. Based on the connectivity information in the graph, we manually linked two contigs with 100 Ns, ultimately obtaining 12 pseudo-chromosomes. Chromosomes were assigned to these 12 scaffolds based on alignment to the 3P Oak genome assembly (Plomion *et al*., 2018). Therefore, the haploid assembly refers to the scaffolded sequences, while the diploid assembly corresponds to the Verkko contigs. We assembled the oak organellar genome using TIPPo v2.1 (Xian *et al*., 2025).

Quality evaluation with Merqury v1.3 indicated a QV of 53 (Rhie *et al*., 2020) with short reads only, and Compleasm assessment showed 99% completeness of the embryophyta_odb10 conserved gene set (Huang and Li, 2023). The same evaluations were also performed on the previous assembly of the Napoleon Oak (Schmid-Siegert *et al*., 2017) for comparison.

Liftoff (Shumate and Salzberg, 2021) was used to project protein-coding genes from the 3P Oak assembly (Plomion *et al*., 2018) onto the Napoleon Oak assembly. Transposable elements were annotated with EDTA v2.0.0 (Ou *et al*., 2019) with parameters --sensitive 1 --anno 1. CpG methylation profiles were identified from HiFi reads by ccsmeth (Ni *et al*., 2023). Satellite repeats were annotated with TRASH (Wlodzimierz, Hong and Henderson, 2023).

To verify the accuracy of the CEN146 array, HiFi reads were aligned to the diploid assembly with Minimap2 (Li, 2018), Winnowmap2 (Jain *et al*., 2022) and VerityMap (Bzikadze, Mikheenko and Pevzner, 2022). StainedGlass (Vollger *et al*., 2022) and pyGenomeTracks (Lopez-Delisle *et al*., 2021) were used to visualize HiFi reads coverage, TE annotation and the similarity of the CEN146 arrays.

CEN146 arrays from homologous chromosomes were aligned separately using Unialigner and Minimap2. Alignment length and mismatch count were analyzed using Unialigner’s built-in script, cigar_histogram.py. To select unique homologous chromosomes, we manually identified two contigs forming a bubble in the assembly graph and aligned them using Minimap2. The specific sequences can be found in Table S7.

### Identification of Somatic Mutations

The Illumina short reads for the upper and lower branches have been published (Schmid-Siegert *et al*., 2017) and were downloaded from NCBI (accession PRJNA327502). Reads were trimmed using fastp v0.23.1 (Chen *et al*., 2018) with the parameters -q 20 -l 75 -w 12 --cut_tail --cut_mean_quality 20.

Trimmed reads were separately aligned to the diploid and haploid assemblies using BWA 0.7.17-r1188 (Li and Durbin, 2009). Potential PCR duplicates were marked using Picard v2.26.7 (*GitHub - broadinstitute/picard: A set of command line tools (in Java) for manipulating high-throughput sequencing (HTS) data and formats such as SAM/BAM/CRAM and VCF*, no date). Minimap2 (Li, 2018) was used to align the diploid and haploid assemblies with asm20; only alignments with MAPQ=60 were kept. Samtools v1.10 stat was used to summarize the alignments.

We extracted the MAPQ and alignment scores for each read from the BAM files with either the haploid or diploid reference genome. If the alignment scores differed, a read was assigned to the reference genome with the higher alignment score. If the alignment scores were the same, the read was assigned to the reference genome with the higher MAPQ value. If the MAPQ values were also identical, the read was assigned to the diploid assembly. Clipped reads were removed.

Reads from the upper and lower branches were used as the mutation and control sample, respectively. When using the diploid assembly as the reference genome, we used bam-readcount v1.0.1 (Khanna *et al*., 2021) with MAPQ≥0 and base quality≥0 for the control sample. Sites with coverage ≥20x, only reference calls, no indels, were collected as control site set A.

For the mutation sample, we applied stringent parameters using bam-readcount v1.0.1 (Khanna *et al*., 2021): MAPQ ≥10, base quality ≥30, site coverage ≤70x. Sites had to have biallelic bases without indels. The alternate allele had to have at least three supporting reads, with support on both strands. Reads with the reference allele had to have an average mismatch ≤0.01. The average mismatch of reads carrying the alternative allele minus the mismatch of reads carrying the reference allele had to be ≤0.01. Only sites overlapping with control site set A were retained.

We used a similar workflow for the haploid assembly as reference, with the following adjustments: control sample, coverage >40x, mutation sample, coverage ≤120x. The somatic mutations identified against both the haploid and diploid assemblies were then combined. For comparison, Sstrelka2 v2.9.10 (Kim *et al*., 2018) was used to detect the somatic mutations using separately either the haploid or diploid assembly.

In diploid assembly, we require a minimum depth of 20× for the control sample. We then quantify regions in the diploid assembly where the sequencing depth is above 20×, resulting in an effective site size of 1.3 Gb.

To identify the position of previously validated SNVs in the new assemblies, we extracted the flanking 100 bp from the previous Napoleon Oak assembly (Schmid-Siegert *et al*., 2017) and aligned the sequences to both our haploid and diploid assemblies with Minimap2 (Li, 2018).

To calculate the allele frequency of the nine somatic SNVs shared between leaves and acorns in the 3P Oak (Plomion *et al*., 2018), we downloaded short reads from NCBI (accession number PRJEB8388). We used the same approach as for the Napoleon Oak to process and align the reads to the assembly. We extracted the allele frequency of the nine SNVs from the bam file.

## Data Availability

Pacbio HiFi sequencing data, haploid and diploid assemblies have been submitted to the ENA under the project accession PRJEB85730. Assemblies in fasta and gfa format and Figure S11 are available through https://doi.org/10.6084/m9.figshare.28397981.v3.

## Code Availability

Scripts are available at

https://github.com/Wenfei-Xian/Somatic_mutation_in_high_heterogyzous_genome

## Acknowledgments

We thank Christian Hardtke for encouragement, intellectual support and discussion. We are especially grateful for the thoughtful feedback from Korbinian Schneeberger. We are grateful to Adrián Contreras, Yueqi Tao, Zhigui Bao, Andrea Movilli and Sebastian Vorbrugg for discussion. This study was supported by the Max Planck Society and the Novozymes Prize of the Novo Nordisk Foundation (D.W.).

## Author Contributions

D.W. designed and supervised the project. P.C-B., F.A.R and P.R. prepared the leaf sample. P.C-B. and W.X. prepared the DNA library. W.X., P.C-B, and F.A.R analyzed the data. W.X., I.B. and D.W. prepared the final manuscript with inputs from all authors.

## Competing Interests

DW holds equity in Computomics, which advises plant breeders. DW also consults for KWS SE, a globally active plant breeder and seed producer. All other authors declare no competing interests.

## Notes

https://doi.org/10.6084/m9.figshare.28397981.v3

