## Supplemental Figures for "Minimizing detection bias of somatic mutations in a highly heterozygous oak genome"

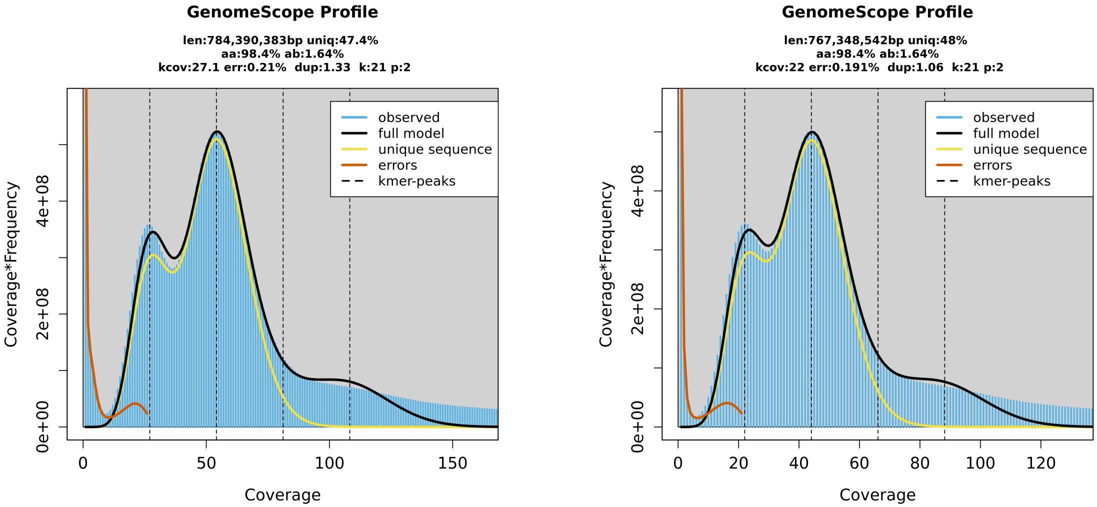


**Figure S1.** Genome survey using GenomeScope2. A. using illumina reads from lower branch. B. using illumina reads from upper branch.

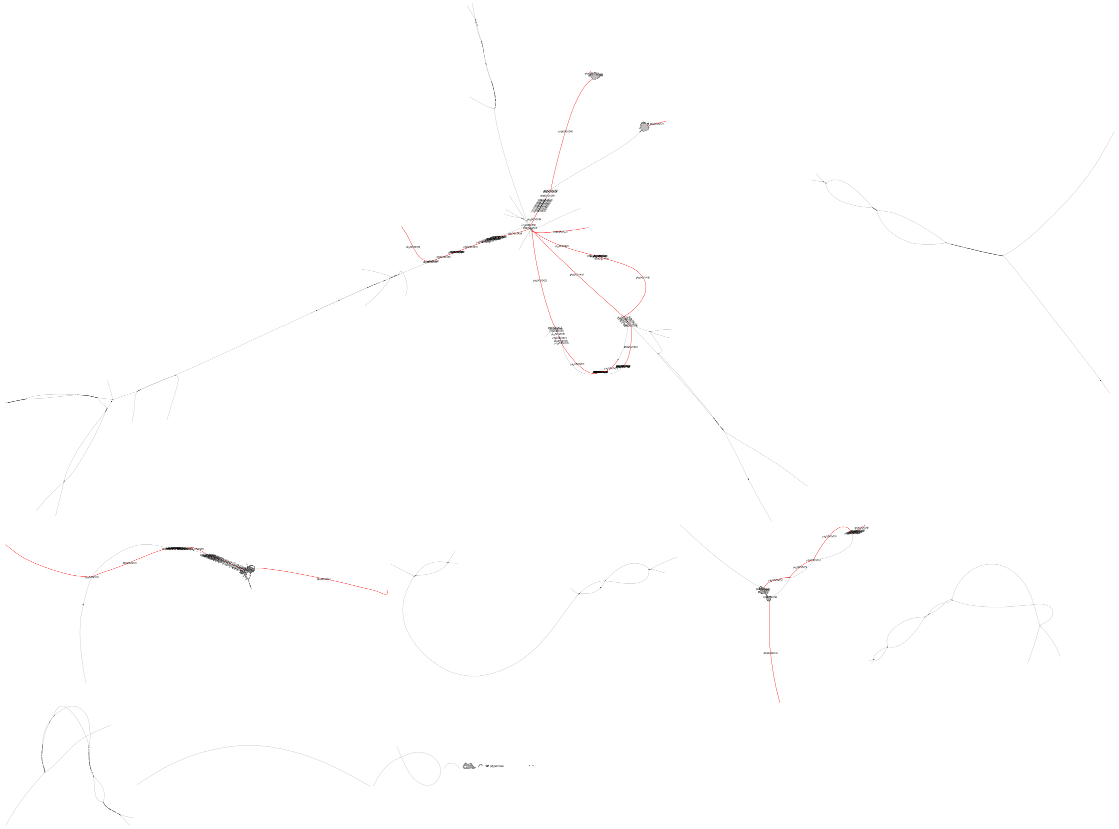

**Figure S2.** Eight long contigs lacking two telomeric repeats in the diploid assembly graph.

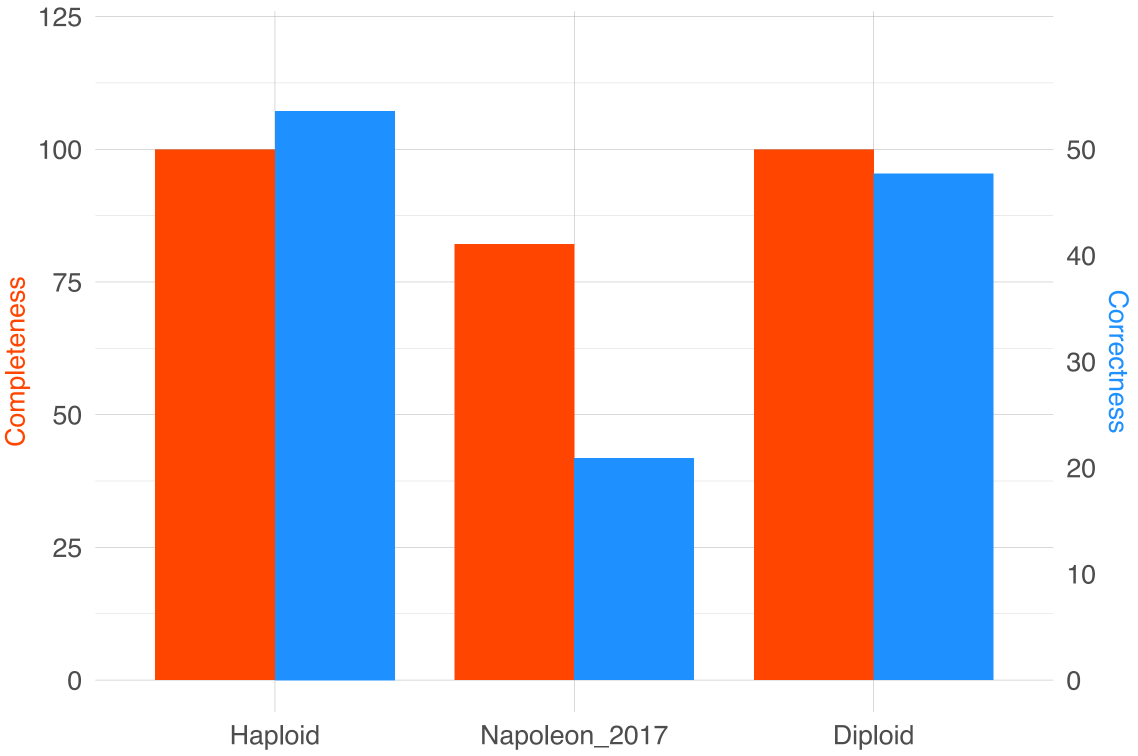


**Figure S3.** Quality evaluation of two assemblies.

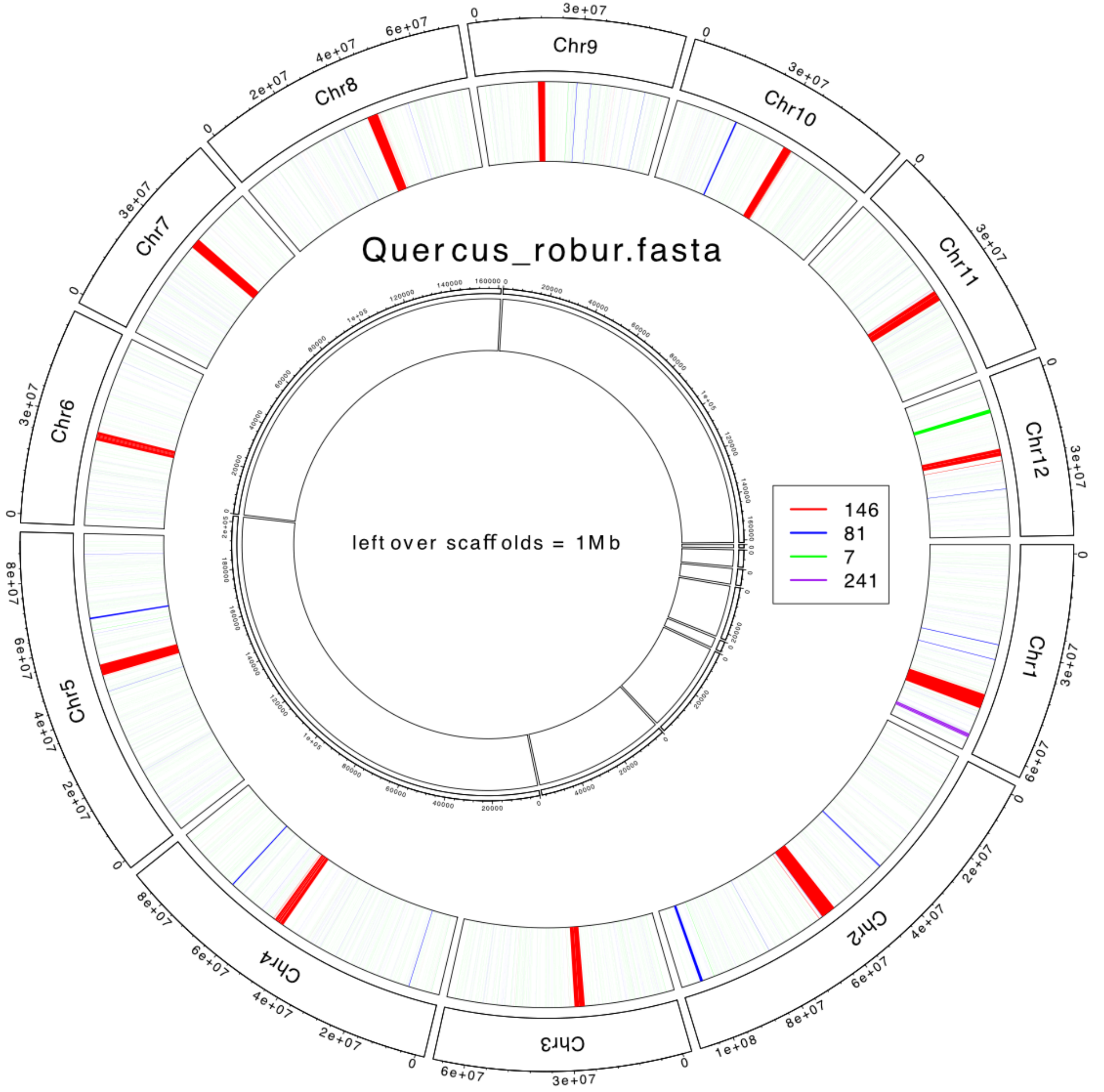


**Figure S4.** Satellite repeats in haploid assembly.

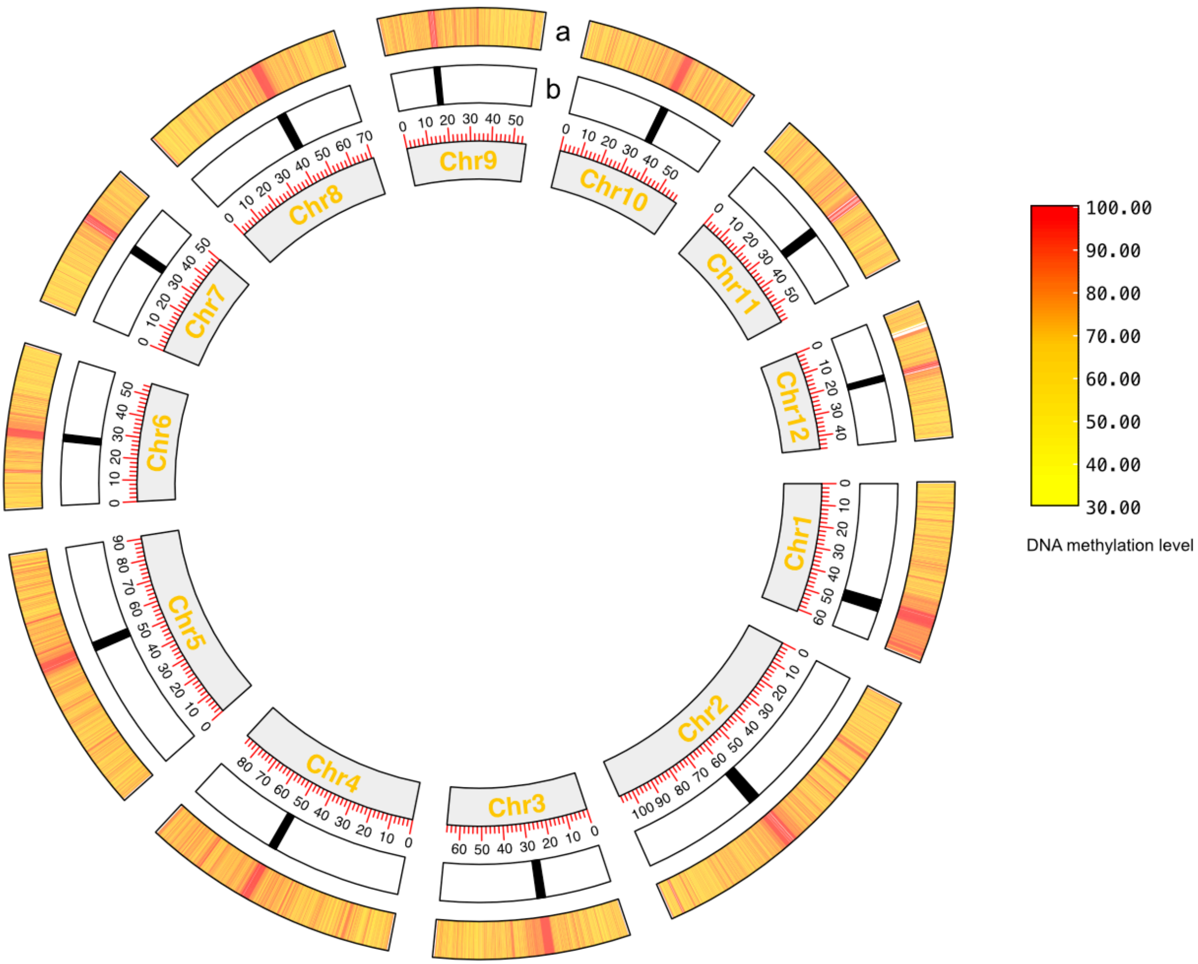


**Figure S5.** Circos plot of CEN146 and DNA methylation. a track indicates the DNA methylation level and b track indicates the location of CEN146.

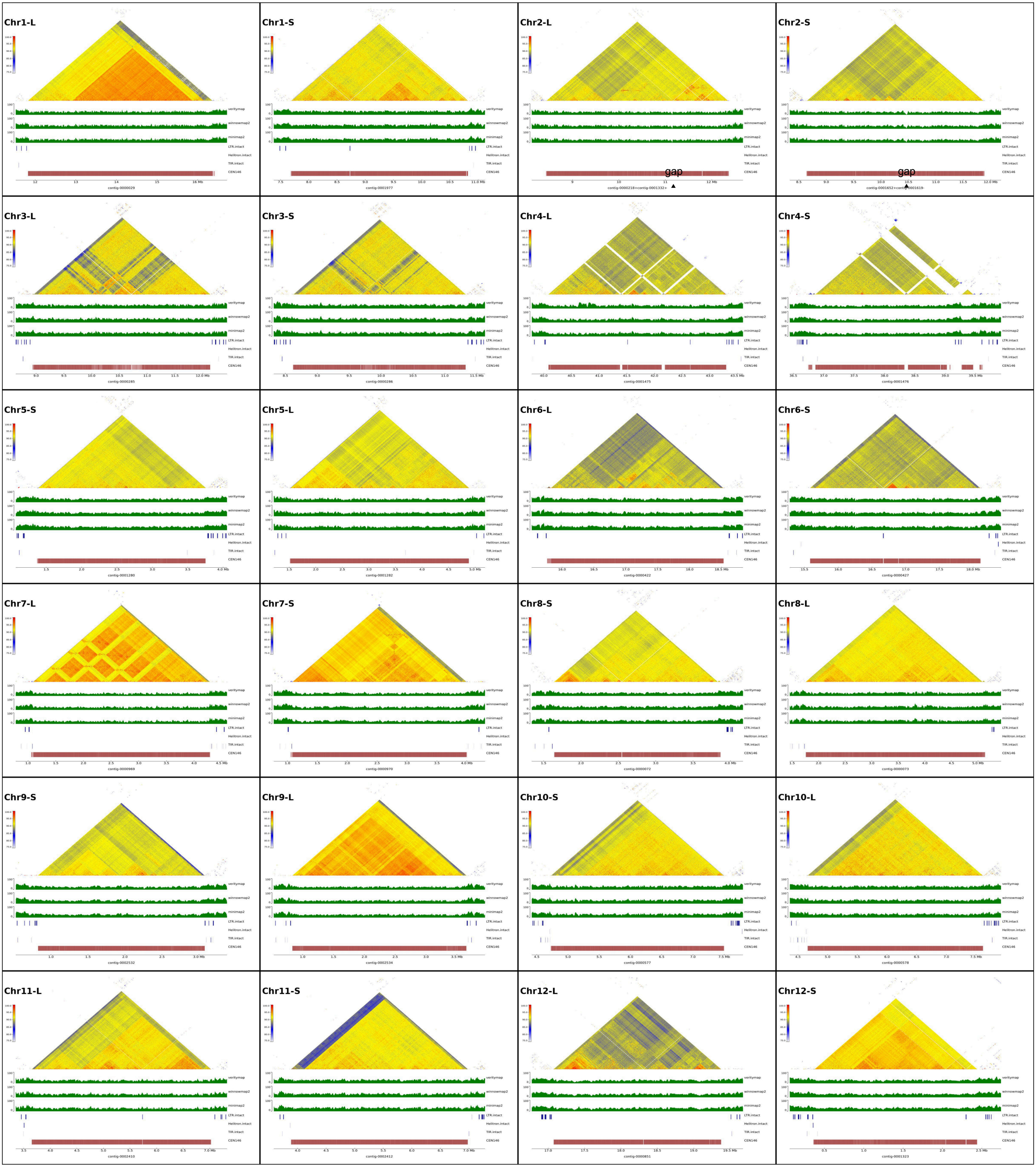


**Figure S6.** HiFi coverage, the present of intact TE and similarity of CEN146 arrays.

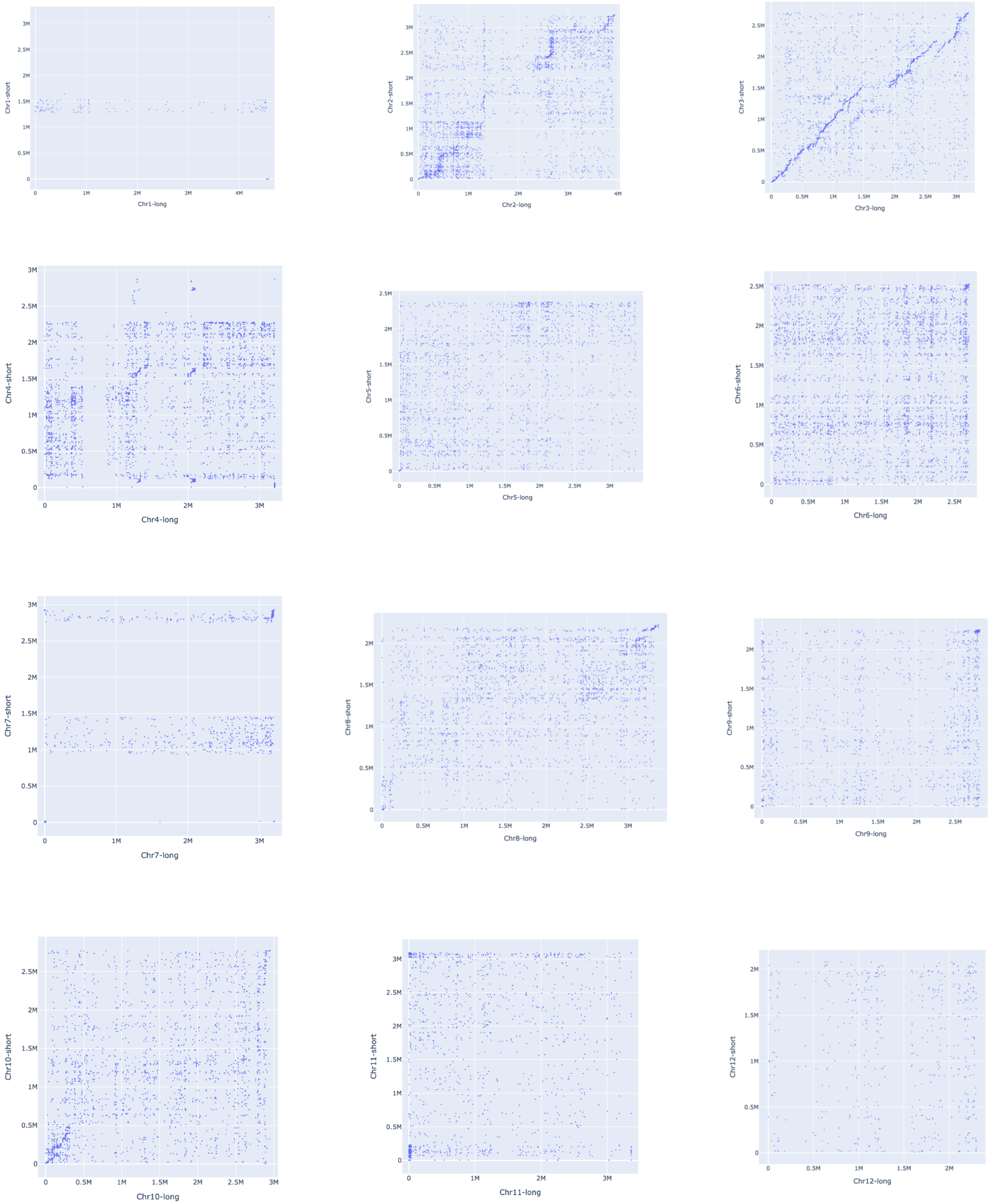


**Figure S7.** Rare dot-plot of CEN146 arrays between homologous chromosome generated by Unialigner.

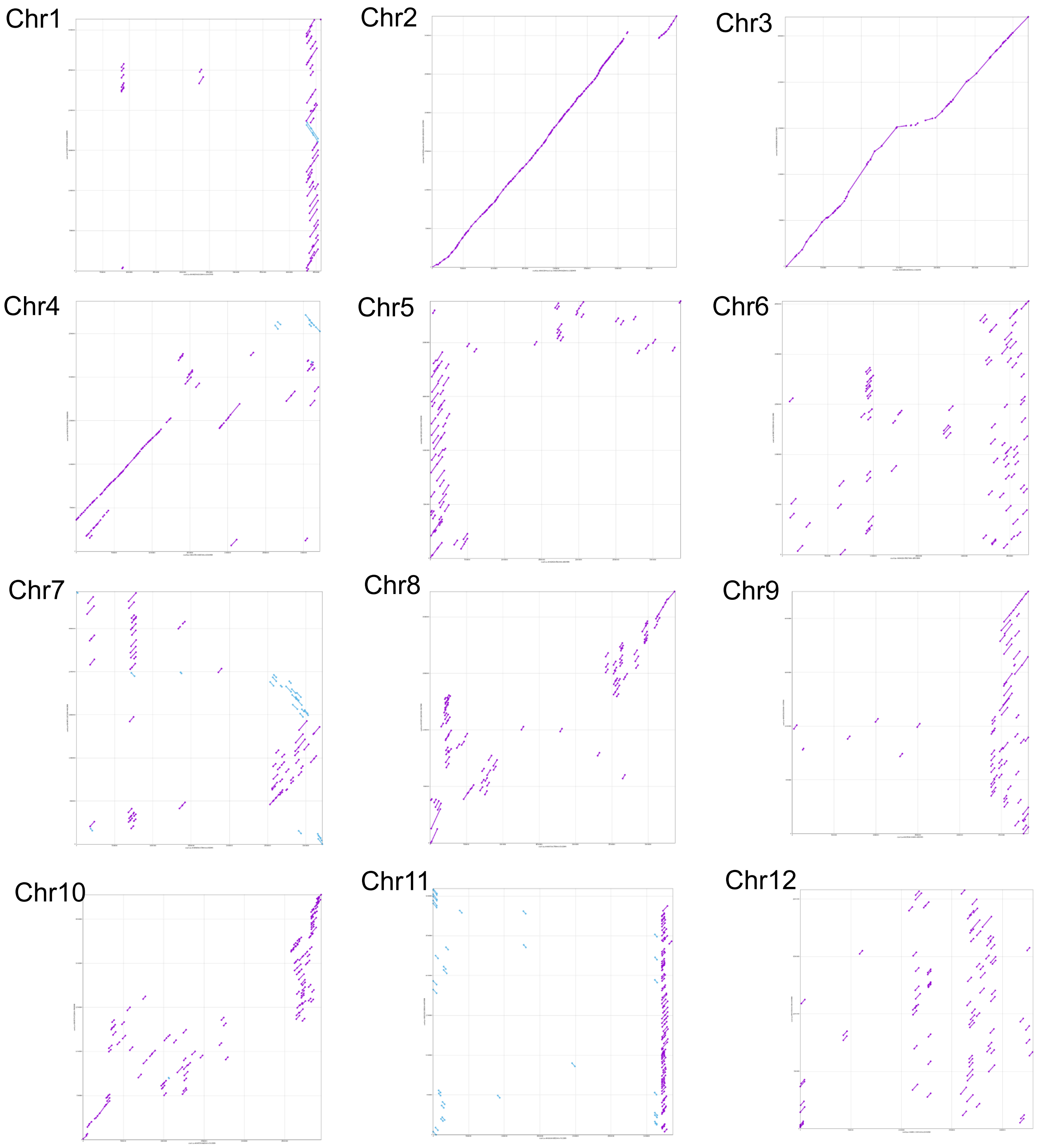


**Figure S8.** Dot-plot of CEN146 arrays between homologous chromosome by Minimap2.

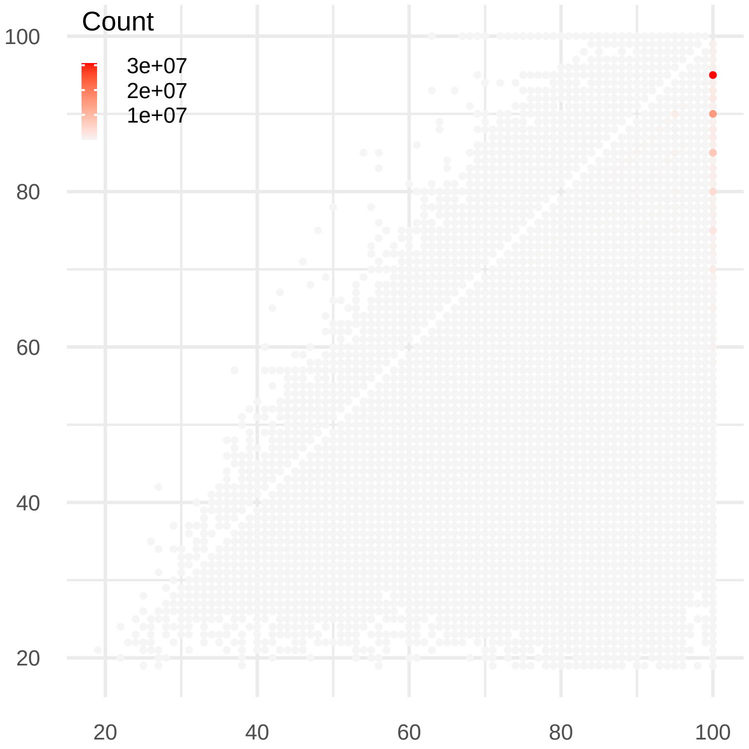


**Figure S9.** Distribution of alignment score in two assemblies as reference. The color intensity of the dots represent the actual number of reads.

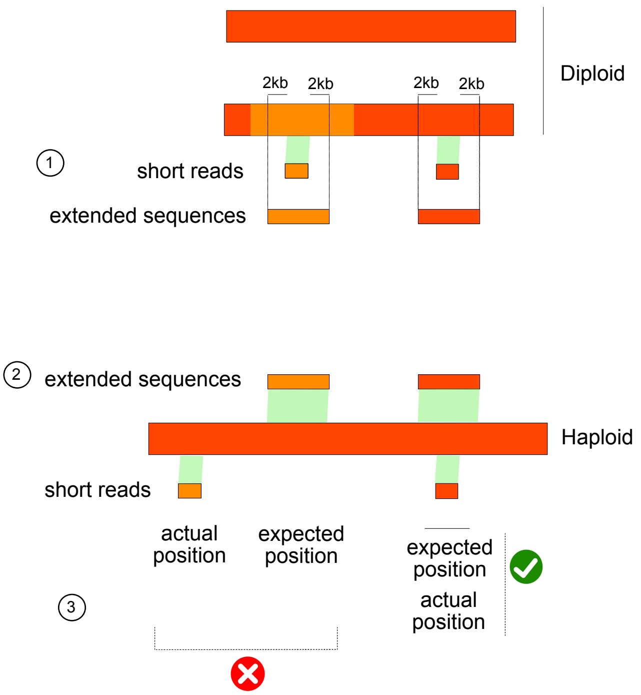


**Figure S10.** Identification of misaligned short reads in the haploid assembly. Step 1: Align short reads to the diploid assembly and extract the flanking 2 kb sequences. Step 2: Align the short reads and the corresponding extended sequences to the haploid assembly. Step 3: Compare the positions of the short reads and the extended sequences.

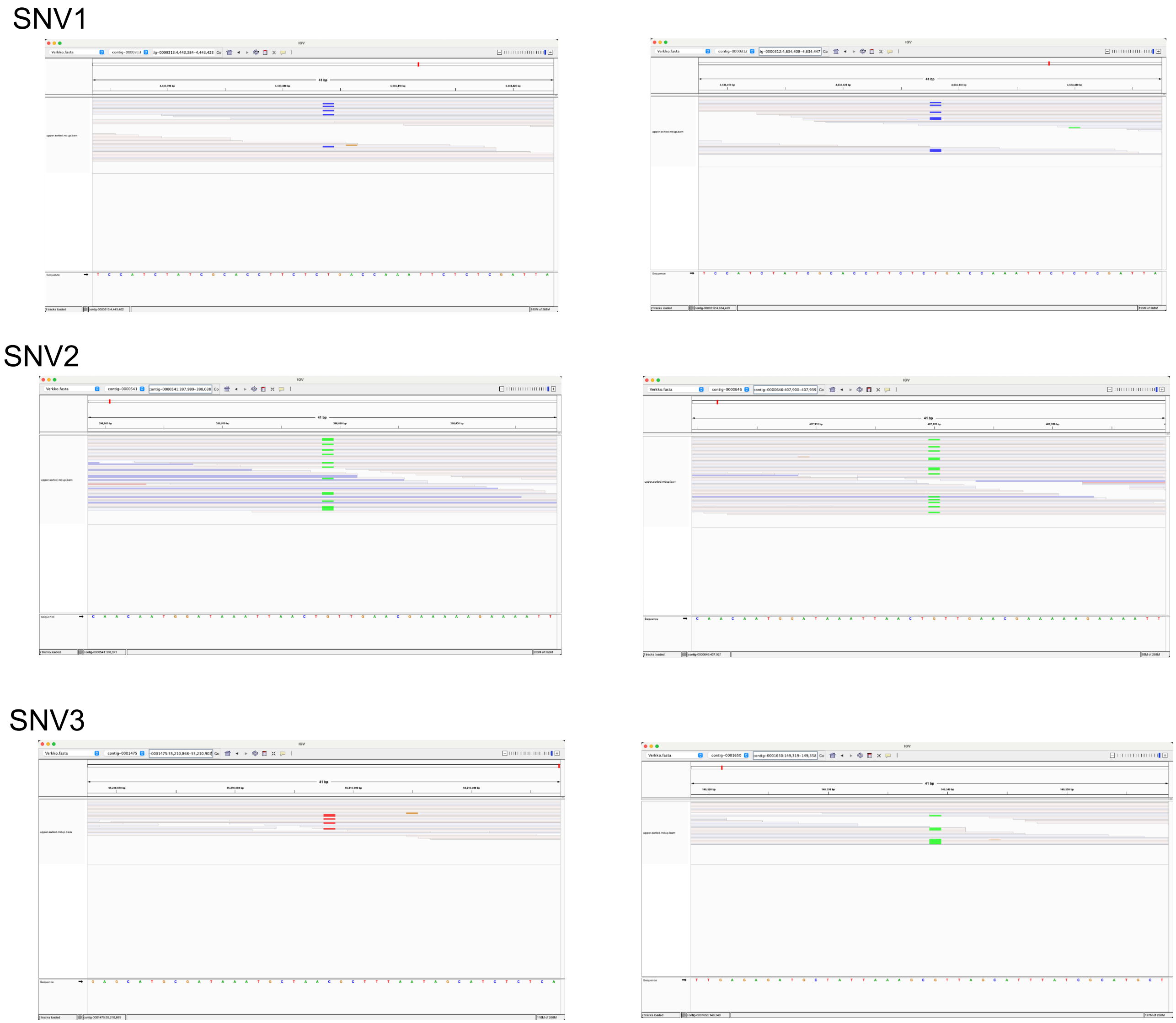


**Figure S12.** Read alignments for the three undetected SNVs using the diploid assembly as the reference.
